# Resting-state connectivity and its association with cognitive performance, educational attainment, and household income in UK Biobank (N = 3,950)

**DOI:** 10.1101/164053

**Authors:** Xueyi Shen, Simon R Cox, Mark J Adams, David M Howard, Stephen M Lawrie, Stuart J Ritchie, Mark E Bastin, Ian J Deary, Andrew M McIntosh, Heather C Whalley

## Abstract

Cognitive ability is an important predictor of lifelong physical and mental well-being and its impairments are associated with many psychiatric disorders. Higher cognitive ability is also associated with greater educational attainment and increased household income. Understanding neural mechanisms underlying cognitive ability is therefore of crucial importance for determining the nature of these associations. In the current study, we examined the spontaneous activity of the brain at rest to investigate its relationships with not only cognitive ability, but also educational attainment and household income. We used a large sample of resting-state neuroimaging data from UK Biobank (N=3,950). Firstly, analysis at the whole-brain level showed that connections involving the default mode network (DMN), fronto-parietal network (FPN) and cingulo-opercular network (CON) were significantly positively associated with levels of cognitive performance assessed by a verbal-numerical reasoning test (standardised β ranged from 0.054 to 0.097). Connections associated with higher levels of cognitive performance were also significantly positively associated with educational attainment (r=0.48, N=4,160) and household income (r=0.38, N=3,793). Further, analysis on the coupling of functional networks showed that better cognitive performance was associated with more positive DMN-CON connections, decreased cross-hemisphere connections between homotopic network in CON and FPN, and stronger CON-FPN connections (absolute β ranged from 0.034 to 0.063). The present study finds that variation in brain resting state functional connectivity associated with individual differences in cognitive ability, largely involving DMN and lateral prefrontal networks. Additionally, we provide further evidence of shared neural associations of cognitive ability, educational attainment, and household income.

## Introduction

General cognitive ability is positively associated with a longer duration of education^1^, better examination performance in school^2^ and better workplace performance^3^. Cognitive ability in youth is also positively associated with higher socioeconomic status in adulthood^3^, and with reduced risk of several mental and physical diseases^3–6^. Thus, cognitive ability is a key trait associated with many educational, social and health outcomes^7^, and therefore identifying the associated neural mechanisms will help better understand the causes of these associations.

Structural and event-related fMRI studies have consistently identified prefrontal brain regions as having the strongest associations with general cognitive ability^8,9^. These regions play a crucial role in executive control^10^ and multisensory integration^11^, and can be assessed using various task-based paradigms^12–14^. However, it has recently been demonstrated that the brain is highly active in the absence of experimental stimuli, i.e. when it is in ‘resting state’. The activity of the brain under resting state is metabolically demanding and topologically efficient; it has been proposed that this actively maintains neural signalling in preparation for quick adaptions^15,16^. Such spontaneous modulations at rest are temporally correlated between distant brain regions, forming the linkage known as functional connectivity.

The spatial patterns of functional connectivity are known as resting state networks (RSN). It is well established that these RSN can be robustly extracted from fMRI data^17^, and they have been consistently verified in several independent cohorts^18^. The RSN approach provides a non-invasive, task-free way of studying such distributed functional dynamics of the brain^19^. In addition to its broad practicability, functional networks found under resting-state are spontaneous, and are therefore free from confounding due to external input^20^. This approach therefore provides the possibility of examining the simultaneous involvement of multiple networks, whose temporal organisation is relevant to cognitive performance that requires various high-level, integrative, cognitive mechanisms^20,21^

RSN involving lateral prefrontal cortex, such as executive control network and frontal-parietal network, have been previously reported to have positive associations with cognitive ability in fMRI studies^22^. Stronger connections involving these networks were also associated with better performance in tasks requiring attention and executive control^23^. This is in line with findings from structural MRI studies that found both decreased fractional anisotropy of the white matter tracts and decreased volume of grey matter in prefrontal cortex were associated with lower fluid intelligence^8,24^. Increasing evidence also suggests that the default mode network (DMN) demonstrates associations with various psychiatric disorders, cognitive test performance^25,26^, and a large number of positive sociodemographic variables^26^. This network shows distinctive metabolic activity compared with networks involving with lateral prefrontal areas^27^, and is also neuroanatomically distinguishable^28^.

The variability of results found in previous studies examining associations between cognitive ability and functional connectivity^22,26,29^ may be due to relatively small sample sizes, often limited to 100 participants or fewer. This limitation is difficult to overcome using meta-analysis, as methods of extracting functional networks may vary considerably between studies. Therefore, there is a need for large-scale studies using a single scanner and consistent methods of estimating the association of RSN activity with consistently-collected social and psychological phenotypes to determine the relationship between resting functional connectivity and cognitive ability.

In the current study, we examined resting-state data from the first release of the UK Biobank imaging project^30,31^. UK Biobank is a large-scale epidemiological cohort where participants from 40 to 75 years old were recruited widely across the United Kingdom^30,32,33^. For the resting-state fMRI (rs-fMRI) data used in the current study, 3,950 subjects had undergone cognitive assessment using a test of verbal-numerical reasoning (VNR; referred to in UK Biobank as a test of “fluid intelligence”). This test had a test-retest reliability of 0.65 between the initial assessment visit in 2006-2010 and the first repeat assessment visit in 2012-2013^34,35^. It also shows a significant genetic correlation with childhood general cognitive ability (r=0.81)^36^.

In addition to the utility of analysing a large sample, the present study benefited from examining the neural associations with educational attainment and household income. The rs-fMRI data were available for educational attainment and household income on samples of 4,160 and 3,793 subjects, respectively. Both education and household income show phenotypic correlations and shared genetic architecture with cognitive ability^35,37^; however, the associations between cognitive ability and these two variables with respect to functional connectivity remain unclear.

In order to address the above issues, we first examined associations between whole-brain connectivity measures and cognitive performance using the VNR task. Second, to compare overlapping networks involved in cognitive performance, educational attainment and household income, we conducted similar whole-brain tests or each trait separately, which were then compared with the results for cognitive performance. Finally, we examined the integrative coupling between these resting state functional network connections using a network-of-interest (NOI) approach, focussing on networks identified by the previous two whole brain analyses.

## Results

### Cognitive performance, educational attainment, and household income

The mean test performance score for the VNR was 6.92 (SD = 2.15). Age and sex both showed significant associations with VNR score (age: β=−0.07, p=3.50×10^−5^, sex: β=0.19, p=3.18×10^−9^ Male=1,Female=0).

In total, 1,801 participants reported having obtained a college/university-level degree (43.29% of the overall sample). The mean age of people with a college/university-level degree was 61.62 (SD=7.49), which was significantly lower than the group without (Mean age=62.65, SD=7.58, t=4.37, p=1.27×10^−5^). Men reported a significantly higher proportion of college degrees (48.80%) than women (39.73%), χ^2^=34.8, df=1, p=3.65×10^−9^. Educational attainment showed positive association with cognitive performance, with age, age^2^ and sex controlled (β=0.457, p<2×10^−16^).

The proportion of people who reported having household income at each level is shown in Figure 1. The income band of £31,000 to £51,999 contained the highest proportion (29.98%) of individuals, and the band >£100,000 contained the lowest proportion (6.06%). Both age and sex showed significant associations with household income (age: β=-0.29, p<2×10^−16^; sex: β=0.20, p=1.04×10^−9^). Higher household income was associated with better cognitive performance (β=0.167, p<2×10^−16^), with age, age^2^, and sex controlled in the model.

**Figure 1.**
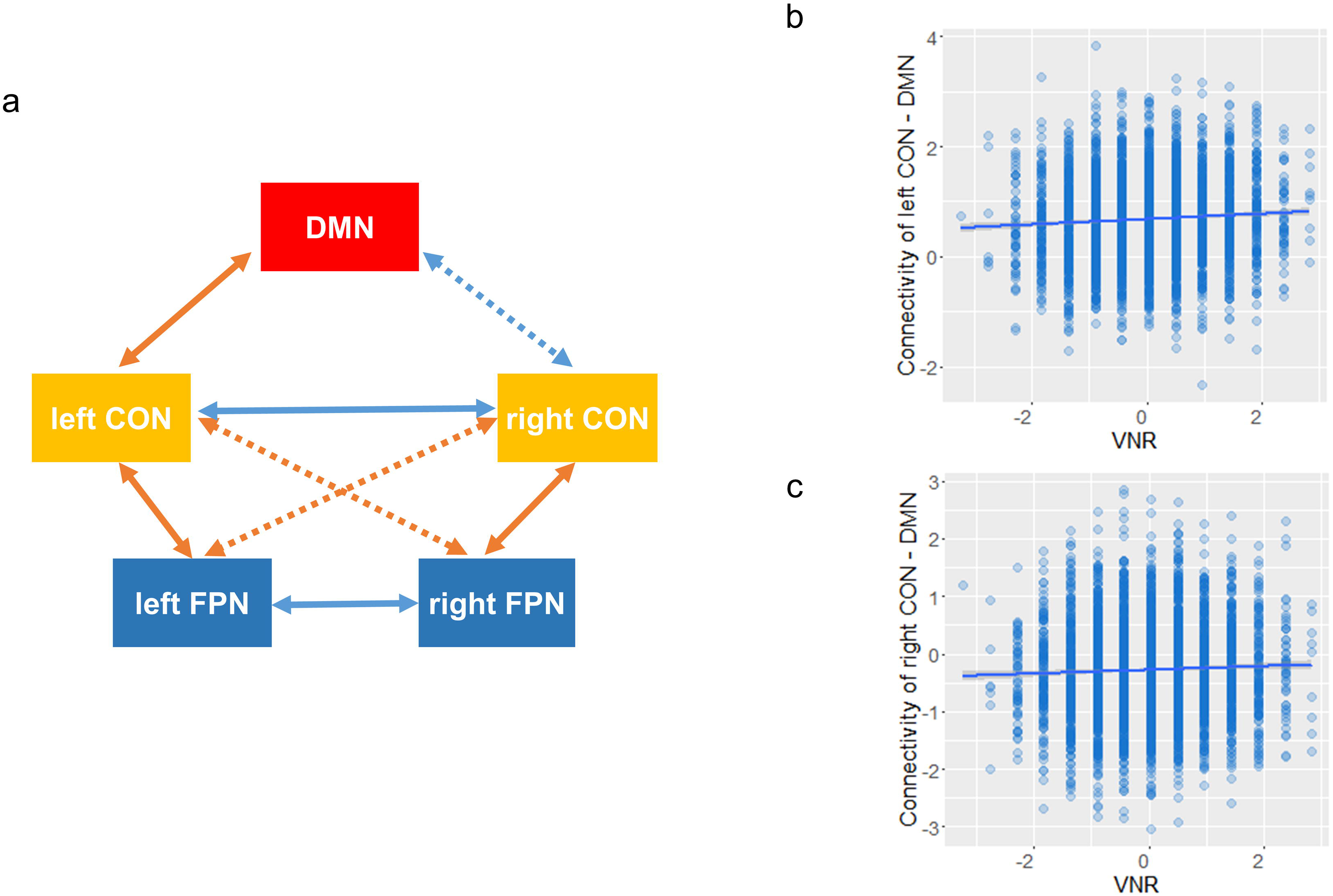
Descriptive statistics of (a) cognitive performance on the verbal-numerical reasoning test; (b) educational attainment (those with [0] and without [1] a college degree; and (c) household income (GBP per annum).

### Whole-brain test of the association of cognitive performance with functional connectivity

A group-ICA was applied to parcellate the whole brain into 55 components, and the pair-wise functional connectivity between the components were estimated using FSLnets (http://fsl.fmrib.ox.ac.uk/fsl/fslwiki/FSLNets). The 55*55 partial correlation matrix was used for whole-brain analysis. To enable clearer interpretation of the results, the values of the connections were transformed into connection strength^26^. This was achieved by multiplying the raw connection values with the signs of their mean value (see Methods).

Better performance in VNR was significantly associated with 26 connections (absolute β ranged from 0.054 to 0.097, all p_corrected_ < 0.05, see Supplementary Table S1). There were 18 significant connections that showed a significant positive association between connection strength and cognitive functioning in VNR; and 8 connections were negatively associated (Supplementary Table S1). The positive connections largely involved the DMN, which includes bilateral posterior cingulate cortex (PCC), bilateral medial prefrontal cortex (PFC) and right temporal-parietal junction (TPJ), see Figure 2. Additional areas of right inferior PFC, dorsal anterior cingulate cortex (ACC) bilateral anterior insula and visual cortex were also involved. Negative associations with cognitive performance included bilateral lateral postcentral gyrus and superior ACC (Figure 2).

**Figure 2.**
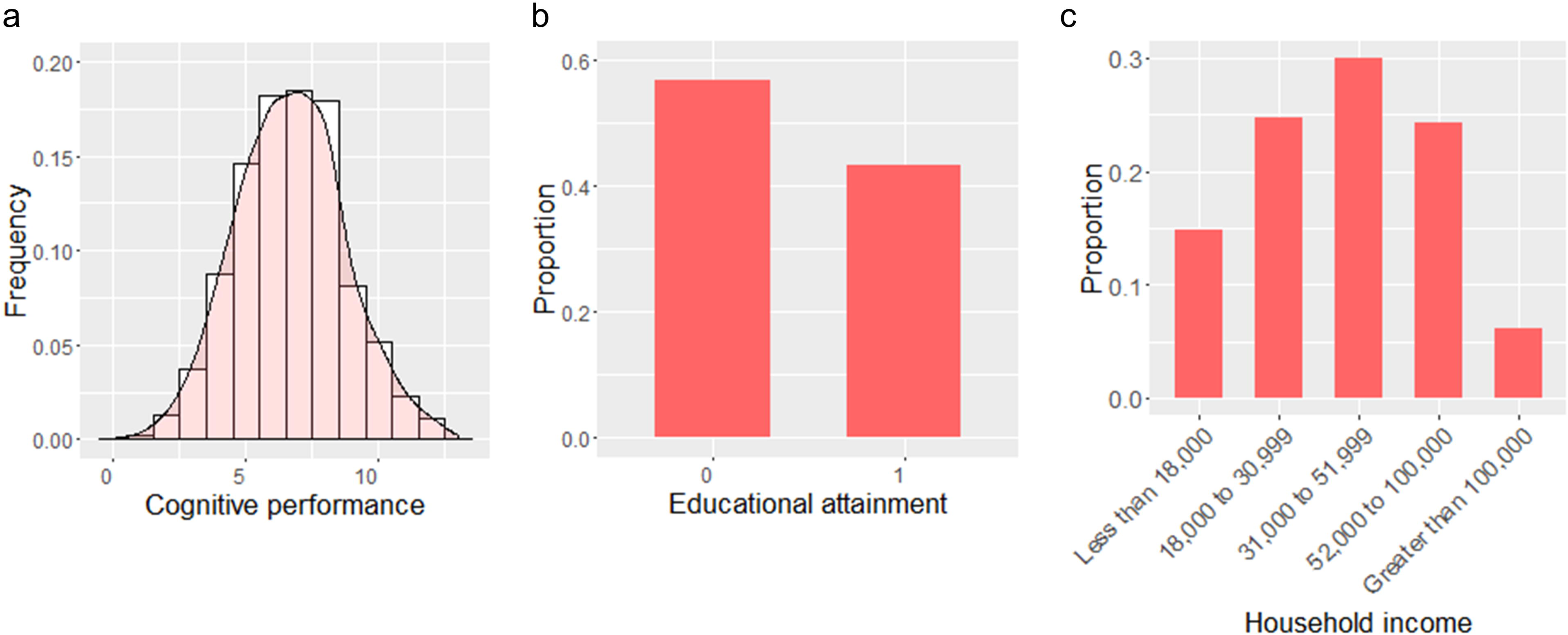
(a) Connections that showed significant associations with cognitive performance. The ICA components were clustered into five categories according to the group-mean full correlation matrix for better illustration and interpretation of the results. This clustering gives a data-driven, gross overview of the structure of the components, consistent with previous studies (ref 26 and 30). The clusters roughly represent the resting state networks (RSNs) of: default mode network (red), extended default mode network and cingulo-opercular network (purple), executive control and attention network (green), visual network (blue) and sensorimotor network (orange). Red lines are the connections where strength was positively associated with cognitive performance; blue lines denote negative associations with cognitive performance. The width of lines indicates the effect sizes of the associations between connection strength and cognitive performance (bigger width indicates a larger absolute effect size). The significant connections were mostly involved in the categories of default mode network, executive control/attention network and cingulo-opercular network. (b) The spatial map of regions involved with connections in (a). The spatial maps for the ICA nodes that involved in the significant connections were multiplied by their effect sizes, then the spatial map in (b) was generated by summing up the weighted maps. To better illustrate the regions involving in significant connections, a threshold of 50% of the highest intensity was applied, so the regions with intensity higher than the threshold would show on the map.

### Whole-brain tests on the association of educational attainment and household income with functional connectivity

There were 33 connections that showed significant associations with educational attainment (absolute β ranged from 0.103 to 0.161, all p_corrected_<0.05, see Supplementary Table S2). Of these, strength of 21 connections were positively associated with higher educational attainment, whereas 12 were negatively associated. The regions involved in positive associations with educational attainment included regions in DMN and dlPFC. A large area of ACC was also involved. Comparatively smaller brain regions were involved in negative associations with educational attainment, which were located in the Inferior part of PCC and lingual gyrus (Figure 3).

**Figure 3.**
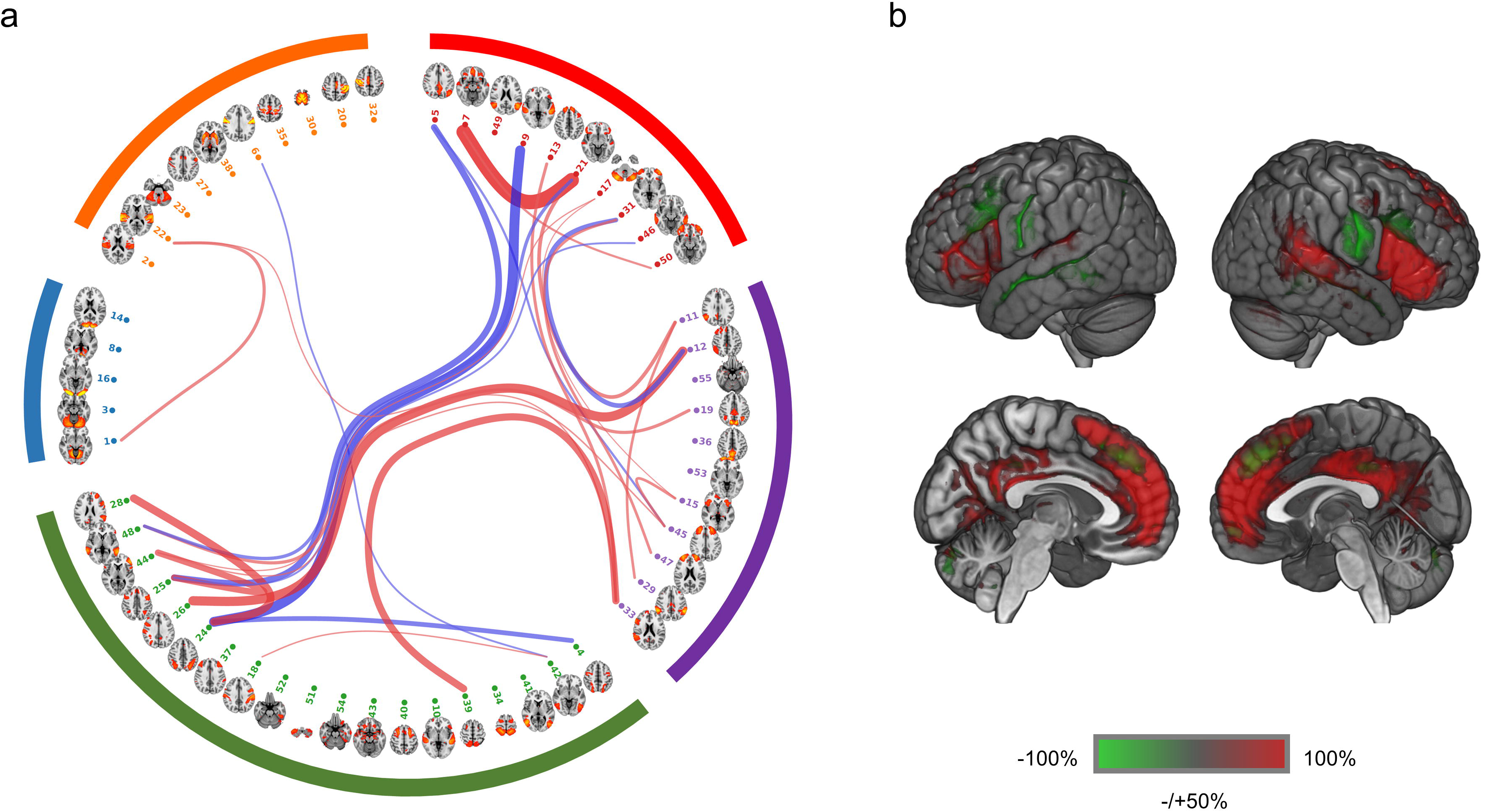
The connections that showed significant associations with educational attainment and household income. Red lines are the connections of which the strength was positively associated with cognitive performance, and the blue lines are the ones having negative associations. The width of lines indicates the effect sizes of the strength of the connections, see the legend of Figure 2. The categorisation of components of brain regions in the circular brain network illustration is identical with Figure 2. Again like Figure 2, A threshold of 50% of the highest value was applied for better illustration of the projection of brain regions on MNI template.

For household income, 15 connections were significant, 11 of which showed positive correlation and 4 showed negative correlation (absolute β ranged from 0.060 to 0.082, all p_corrected_ < 0.05, Supplementary Table S3). The regions of the positive correlations again fell in similar regions as in tests of educational attainment and cognitive performance, which included PCC, medial PFC, ventral lateral PFC and dorsal lateral PFC (Figure 3). The area that showed negative correlation was smaller, which mainly included superior temporal lobe. Full lists of regions of the maps for the above results are presented in Supplementary Table S4.

The spatial maps for the results of cognitive performance in VNR, educational attainment, and household income overlapped substantially (Figures 2 and 3). By performing correlation analysis at the standardised effect sizes of whole brain (see Methods, Statistical methods), we found a correlation of r=0.47 (df=1,483, p<2×10^−16^) between the global effect sizes for cognitive performance and educational attainment. The correlation between the effect sizes of cognitive performance and household income was r=0.38 (df=1,483, p<2×10^−16^) (Figure 4).

**Figure 4.**
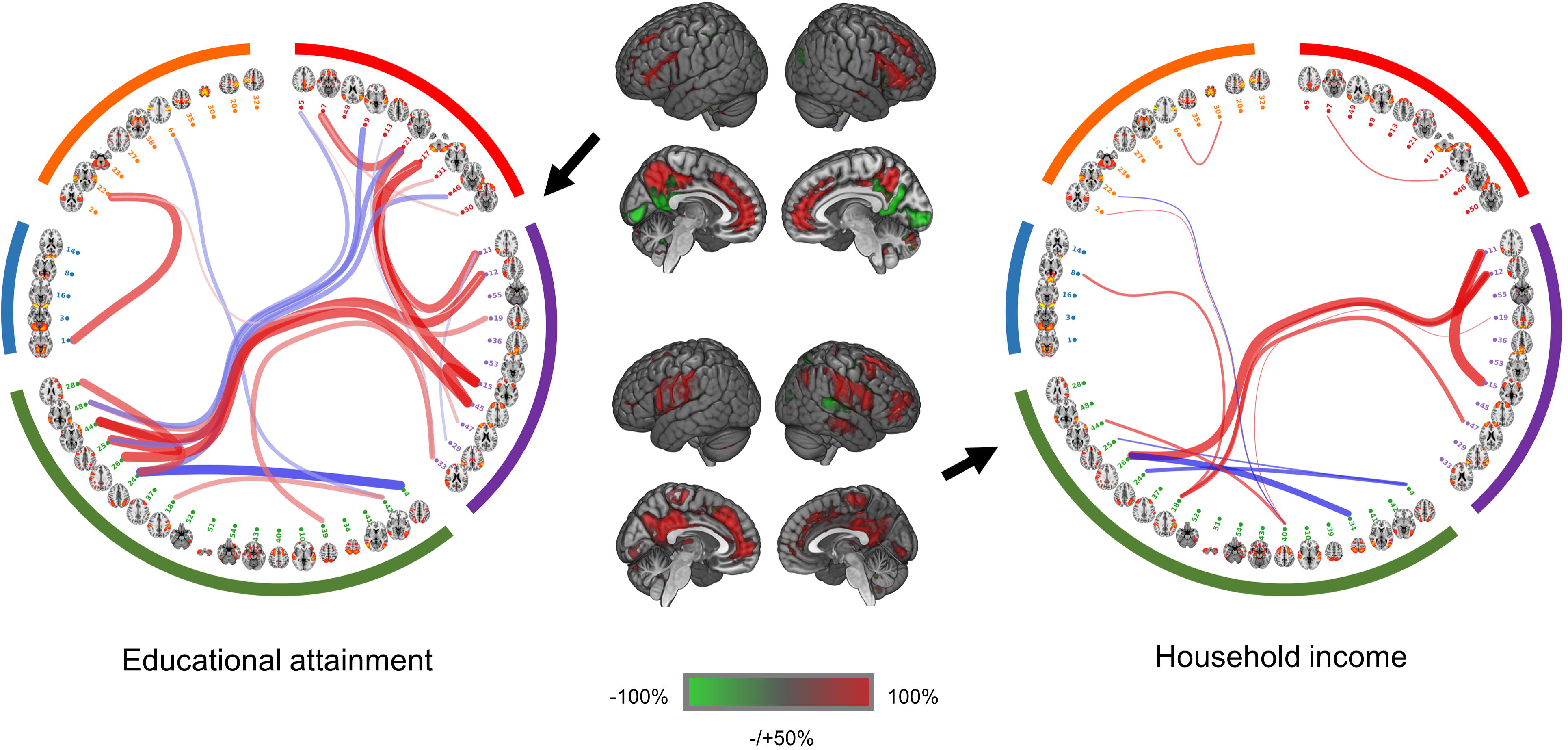
Correlations of the effect sizes of (a) cognitive performance and educational attainment and (b) cognitive performance and household income on whole-brain connections using 55*55 partial correlation matrix as the proxy. Regression line with 95% confident intervals (shaded) are shown.

### Network-of-interest (NOI) test on VNR, educational attainment, and household income

The whole-brain tests showed that the connections associated with cognitive performance in VNR, educational attainment and household income were predominantly located within the DMN (covering medial PFC, PCC and TPJ), cingulo-opecular network (CON, covering ventral lateral PFC, and dorsal ACC) and fronto-parietal network (FPN, covering dorsal lateral PFC and posterior parietal cortex). Therefore, DMN, CON, and FPN were selected as NOI from another group-ICA of lower resolution so these networks could be fully extracted (see Methods). The pairwise between-network coupling of these five networks (DMN was unilateral, and CON and FPN were separately extracted on each hemisphere) was tested to determine their association with cognitive performance, educational attainment, and/or household income. The spatial maps for the above components can be viewed in Figure 5. The valence and values for the coupling of the above NOI were shown in Table 1. Similar with the analyses at whole-brain connectivity, the values of the connections were transformed into connection strength before they were fed into the model.

**Table 1.** The significant associations between the couplings of networks of interest and cognitive performance (verbal-numerical reasoning) and educational attainment. The values of connections were transformed into strength before conducting the analyses, by multiplying the connection values with the signs of their means. This approach was consistent with ref 28. Mean values and their 95% confident intervals of connections reported here are the values before being transformed into strength.

| **Verbal-Numerical Reasoning** |  |  |  |  |  |  |  |  |  |
| --- | --- | --- | --- | --- | --- | --- | --- | --- | --- |
| Type | Connections | Beta | Standard error | t.value | p | pcorrected | Mean value of connection | 95% confident interval of value of connection |  |
| inter-hemisphere | left FPN - right FPN | -0.040 | 0.016 | -2.493 | 1.27E-02 | 0.018 | 1.156 | 1.127 | 1.185 |
|  | right CON - left CON | -0.063 | 0.016 | -3.923 | 8.89E-05 | 6.67E-04 | 0.379 | 0.356 | 0.402 |
| CON - FPN | left CON - right FPN | 0.034 | 0.016 | -2.106 | 3.52E-02 | 0.044 | -1.359 | -1.387 | -1.330 |
|  | right CON - left FPN | 0.043 | 0.016 | -2.714 | 6.68E-03 | 0.011 | -2.088 | -2.122 | -2.054 |
|  | left CON - left FPN | 0.044 | 0.016 | 2.732 | 6.33E-03 | 0.011 | 1.043 | 1.018 | 1.067 |
|  | right CON - right FPN | 0.051 | 0.016 | 3.200 | 1.38E-03 | 0.005 | 0.648 | 0.620 | 0.676 |
| DMN-related | left CON - DMN | 0.061 | 0.016 | 3.824 | 1.33E-04 | 6.67E-04 | 0.675 | 0.652 | 0.698 |
|  | right CON - DMN | -0.045 | 0.016 | 2.797 | 5.18E-03 | 0.011 | -0.275 | -0.300 | -0.250 |
| **Educational attainment** |  |  |  |  |  |  |  |  |  |
| Type | Connections | Beta | Standard error | t.value | p | pcorrected | Mean value of connection | 95% CI of value of connection |  |
| CON - FPN | right CON - right FPN | 0.086 | 0.031 | 2.736 | 6.24E-03 | 0.021 | 0.648 | 0.620 | 0.676 |
| DMN-related | right FPN - DMN | 0.104 | 0.031 | -3.335 | 8.59E-04 | 0.004 | -0.710 | -0.738 | -0.682 |
|  | right CON - DMN | -0.149 | 0.031 | 4.761 | 1.99E-06 | 1.99E-05 | -0.275 | -0.300 | -0.250 |

Educational attainment

**Figure 5.**
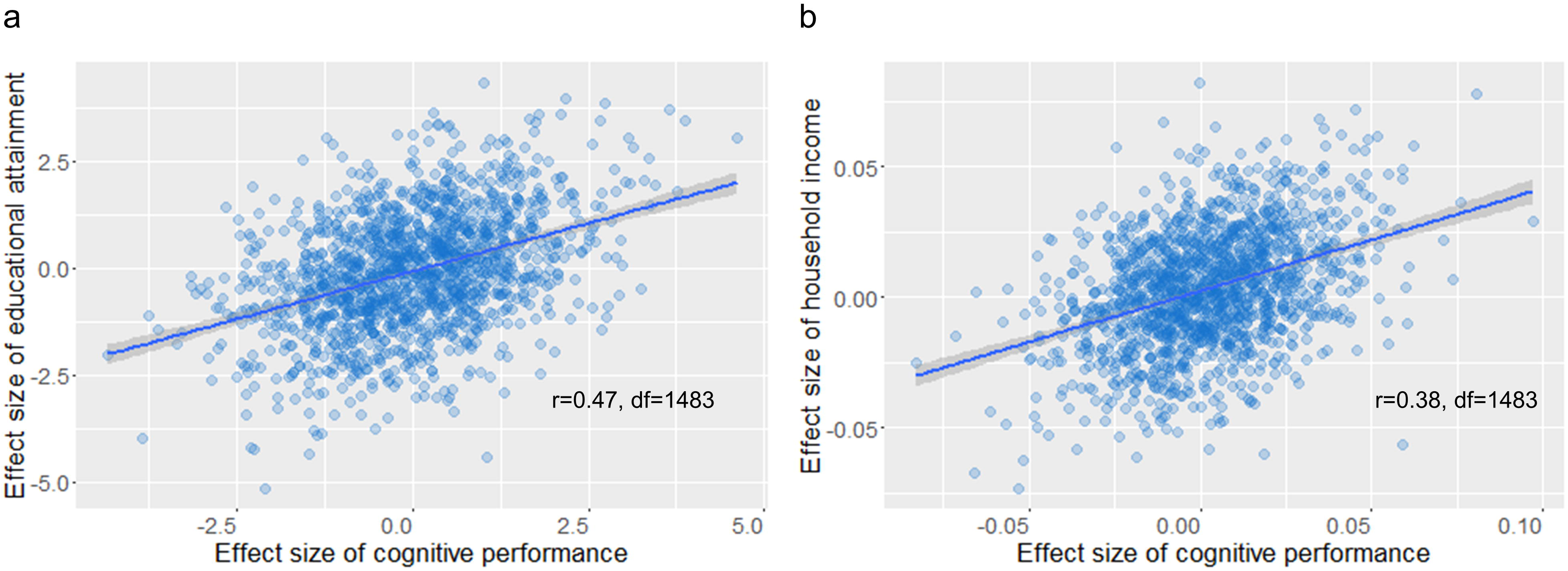
(a) Five intrinsic functional networks selected from the 21 components generated by low-dimension ICA (see Methods, Imaging data). Component 1 was identified as the default mode network (DMN, red). Component 13 and 21 were left and right cingulo-opercular network (CON) respectively (yellow). And finally, component 5 and 6 were identified as right and left fronto-parietal network (FPN, blue). (b) The mean values of couplings of networks of interest. The values are standardised temporal correlation coefficient between network of interest. A higher absolute value indicates a higher strength, and the sign indicates the directionality of the connection. A negative value means an anti-correlated connection, whilst a positive value indicates a positive connection. Mean values and 95% confident intervals of the connections can be viewed in Table 1.

There were 8 couplings of functional networks significantly associated with VNR performance out of 10 connections tested. For educational attainment, 3 connections were significant, and none was found significantly associated with household income.

For the connections between DMN and networks involving with lateral PFC, better VNR performance was associated with more positive connections between DMN and bilateral CON (stronger positive connection between DMN and left CON: β=0.061, p_corrected_=6.7×10^−3^; weaker negative connection of DMN with right CON: β=−0.045, p_corrected_=0.011).

On the other hand, greater strength of connections within the networks involving with lateral PFC were significantly associated with better cognitive performance. Stronger CON-FPN coupling was also associated with higher VNR score. In the same hemisphere, people with better cognitive performance showed higher positive CON-FPN connections (left CON-left FPN: β=0.044, p_corrected_=0.011; right CON-right FPN: β=0.051, p_corrected_=0.005), whilst across hemispheres, negative CON-FPN connections were higher (left CON-right FPN: β=0.034, p_corrected_=0.044; right CON-left FPN: β=0.043, p_corrected_=0.011). Finally, higher VNR scores were associated with weaker cross-hemisphere connections between homotopic network components (left-right FPN:β=−0.040, p_corrected_=0.018. left-right CON: β=−0.063, p_corrected_=6.7×10^−4^). The above results are presented in Table 1 and Figure 6.

**Figure 6.**
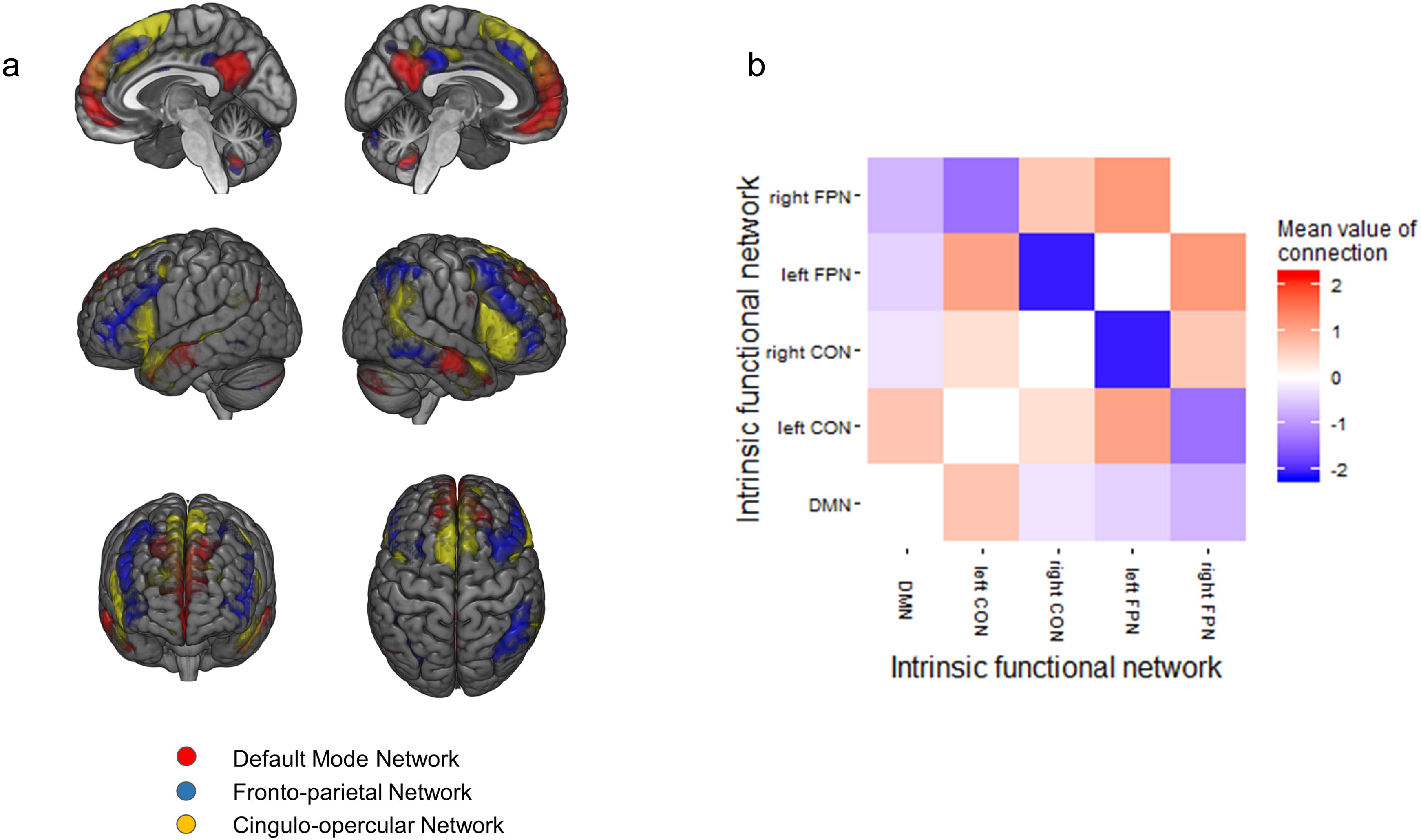
(a) Significant network couplings associated with cognitive performance in verbal-numerical reasoning (absolute β ranged from 0.034 to 0.063, all effect sizes of the significant connections are reported in Table 1). An orange arrow means positive association between cognitive ability with the absolute strength of a connection, whilst a blue arrow indicates decreased absolute strength of a connection with better cognitive performance. Solid arrows are positive connections and dashed ones are negative. (b) and (c) represent the association of cognitive performance in verbal-numerical reasoning and the connection between left/right CON (β = 0.061 and −0.045 respectively for left/right CON) and DMN (β=−0.045). Y-axis represent the normalised correlation coefficient between temporal modulations of networks. Better cognitive performance was associated with more positive connections between DMN and bilateral CON. The spatial maps of the functional networks can be found in Figure 5.

Educational attainment and household income had generally smaller associations with network coupling, and fewer significant connections were found. People with higher educational attainment showed stronger connection between DMN and right FPN (β=0.104, p_corrected_=0.004) and lower connection between DMN and right CON (β=−0.149, p_corrected_=1.99×10^−5^). A stronger positive connection between right FPN and CON was associated with better educational attainment (β=0.086, p_corrected_=6.24×10^−3^). No significant association between household income and the coupling of networks was found (all p_corrected_>0.124).

## Discussion

In the present study, we utilized a large population-based sample of ∼4,000 participants and found that strength of connections involved with DMN regions, anterior insula, and lateral prefrontal areas were positively associated with performance in a verbal-numerical reasoning test. The brain regions associated with cognitive performance also overlapped with those related to educational attainment and household income. For cognitive performance in particular, better cognitive functioning was marked by a more strongly positive DMN-CON connection, weaker cross-hemisphere connections of the left-right CON and left-right FPN, and stronger CON-FPN connections.

We used a large sample and provided evidence that, in addition to the broadly suggested idea of lateral PFC playing a crucial role in cognitive processing, DMN was also associated with cognitive performance (β of connections positively associated with cognitive ability ranged from 0.054 to 0.097) ^8,24,38^. Previous studies showed that DMN serves as a hub for the whole brain^28^. In comparison with other functional networks, DMN showed a higher metabolic rate in resting-state^27^, stronger connections with the rest of the whole brain in both task-free and task-engaging situations^39^, and a key role in maintaining basic levels of wakefulness/alertness in the brain^40^. Higher efficiency within the DMN was reported to be associated with various cognitive functions, including memory^41^, theory of mind^42^, working memory^43^, and performance in general intelligence tests^44^. The high-level cognitive abilities mentioned above often involve the activity of multiple, spatially distant brain regions^41,45^. Therefore the DMN, as a communicative hub, contributes to functional efficiency over the whole brain^44^, potentially producing better integration and cooperation in core regions that are important for cognitive tasks.

Additionally, the present study tested the coupling between networks of interest. More positive DMN-CON coupling was associated with better cognitive ability (absolute β>0.045). In addition to the well-recognised task-positive lateral prefrontal cortex (therefore anti-correlated with the DMN), our findings in this large single-scanner sample lend substantial credence to increasing evidence that the CON itself^18,46^, and its positive coupling with the DMN (in both resting-state^47^ and event-related studies^48^) is highly pertinent for important aspects of cognitive performance. The role of the CON was related with maintaining task-engaging status^18,49^ and flexibly switching between the DMN and central executive network based on experimental context^21,50^. The experimental context in which CON and DMN were found to be simultaneously activated was often about goal-directed cognition^21^, which involves self-driven retrieval of memory or learned experience and self-regulatory planning^29^. As the DMN is associated with self-referential processing^28^ and self-driven cognition like retrieval of personal experience^51^ and planning^29,52^, positive coupling of the CON and DMN may indicate recruitment of self-referential and goal-oriented activity. Therefore successful DMN-CON coupling may be useful in maintaining internal mechanisms that support cognitive processing and long-term learning^21^.

The coupling of networks involving lateral PFC showed that better cognitive performance was associated with stronger CON-FPN connections (absolute β>0.034). This result is consistent with previous structural and functional findings that support the key role of prefrontal areas on cognitive performance^8,53^. We also found that better cognitive performance was related to between-hemisphere dissociation within networks (absolute β>0.040).Whereas this is the first time to our knowledge that this has been examined in a study of large sample, such reduced structural coupling between left and right lateral PFC has been observed in schizophrenic patients with impaired cognitive performance^54^. More lateralisation of the brain is associated with better cognitive performance^55,56^, whereas, less lateralisation, especially in prefrontal cortex, is related with reduced specialisation of brain functions across hemispheres, therefore the advantageous de-coupling we report here potentially denotes increased brain efficiency^55,57^.

The whole-brain connection map for cognitive performance overlaps substantially with those from educational attainment and household income. Further analyses showed that there were global correlations of cognitive ability with educational attainment (r=0.47) and with household income (r=0.38).GWAS studies found that cognitive performance and educational attainment share a similar genetic architecture (r=0.906) ^1,36^. Early life intelligence (relatively stable across the life-course^58,59^) and educational duration show partially overlapping associations with some structural brain measures in older age^60^. Taken together, one interpretation of these data is that the functional hallmarks of a more ‘intelligent’ and better educated brain are related to income by virtue of these temporally preceding factors. It could equally be the case that income confers additional lifestyle benefits that also influence these cerebral characteristics; the causal direction that gives rise to the highly overlapping functional connectivity reported here would be more adequately addressed with longitudinal multi-modal data.

A limitation for the current study is that the verbal-numerical reasoning test, as a brief measure, may not confer the same level of reflection on general cognitive ability as other longer, in depth general cognitive measures. However, as previous studies found that verbal-numerical reasoning shared significant genetic and phenotypic correlation with the latent component of general cognitive performance^35,36^, it therefore confers adequate representativeness of general cognitive ability. A notable strength of the present study is that we used a large sample, providing compelling evidence that both dorsal prefrontal areas and DMN were associated with cognitive ability, educational attainment and household income. To disentangle how multiple networks involved in the cognitive ability, we examined functional connectivity by estimating connections between brain components derived in two different resolutions, giving us another strength of studying both the connections over the whole brain and the coupling of bulk intrinsic functional networks within a single dataset. Finally, in addition to visual checking of overlapping regions of the significant connections, we statistically compared the functional connectivity associated with cognitive ability, educational attainment and household income over the whole brain, giving a magnitude of neural associations among them.

## Conclusion

The present study used a large, population-based sample, who provided multi-dimensional rs-fMRI data, and found substantial evidence for functional neural associations cognitive ability (verbal-numerical reasoning) both in whole-brain dynamics and the coupling of intrinsic functional networks. The findings also characterised the degree of rs-fMRI overlap between cognitive ability and educational and socioeconomic level, providing evidence of the overlapping biological associations on the neurological level.

## Methods

### Participants

The study was approved by the National Health Service (NHS) Research Ethics Service (reference: 11/NW/0382), and by the UK Biobank Access Committee (Project #4844). Written consent was obtained from all participants.

In total, 4,162 participants undertook a rs-fMRI assessment and passed the quality check undertaken by UK Biobank (http://www.fmrib.ox.ac.uk/ukbiobank/nnpaper/IDPinfo.txt) (Mean Age=62.20+/−7.56 years, Male=47.48%).

### Imaging data

We used the network matrices from the IDPs (imaging-derived phenotypes) which were processed by the UK Biobank imaging project team^30^. All the processing of resting-state data described in this section was performed by the UK Biobank team, including acquisition and pre-processing of resting-state data and estimation of functional connectivity. Quality check was conducted by UK Biobank following the standard protocol^61^. The detailed methods of the UK Biobank imaging processing can be found in a previous protocol paper^30^. For clarification, these processes are described briefly below.

All imaging data were obtained on a Siemens Skyra 3.0 T scanner (Siemens Medical Solutions, Germany). The fMRI scans employed a single-shot gradient-echo echo-planar imaging (EPI) sequence with a voxel resolution of 2.4 mm ^3^. Full details of data acquisition are outlined elsewhere: http://biobank.ctsu.ox.ac.uk/crystal/refer.cgi?id=2367.

Data pre-processing, group-ICA parcellation and connectivity estimation were carried out using FSL packages (http://biobank.ctsu.ox.ac.uk/crystal/refer.cgi?id=1977) by UK Biobank. Briefly, pre-processing included motion correction, grand-mean intensity normalisation, high-pass temporal filtering, EPI unwarping, gradient distortion correction unwarping and removal of structured artefacts^30^.

Group-ICA were then performed on the full sample of 4,162 people whose resting-state data was collected and pre-processed. The analyses were conducted using FSL’s MELODIC tool (https://fsl.fmrib.ox.ac.uk/fsl/fslwiki/MELODIC), and two different ICAs were performed with the dimensionality (*D*) set as 100 and 25. The *D* determines the number of distinct ICA components. The dimensionality of *D*=100 infers a larger number and therefore smaller functional components of the whole brain, whilst setting *D*=25 results in larger functional networks that can be extracted as a single component^26,30^. After the group-ICA, components that were white matter-, motion-, and cardiac-related were considered as noise and therefore discarded; this resulted in 55 components in 100-*D* ICA and 21 components in 25-*D* ICA that remained for further analysis. The maps of both ICAs can be seen at: http://www.fmrib.ox.ac.uk/datasets/ukbiobank/index.html.

Finally, connections between pairs of ICA components for each subject were estimated. Two types of matrices were calculated using the FSLNets toolbox: http://fsl.fmrib.ox.ac.uk/fsl/fslwiki/FSLNets. The first type was the full correlation matrix. This was achieved by estimating temporal component-to-component correlation. The estimation of a connection provided by this approach confounds both direct and indirect connectivity via other components. To estimate direct connectivity better, a partial correlation matrix was generated as a second type of proxy, which controlled for the strength of other connections, allowing us to estimate the strength of a connection over indirect paths consisting of other connections. The estimation of partial correlation was conducted with an L2 regularization applied (rho=0.5 for Ridge Regression option in FSLnets). The values of connections were normalised.The parameters for this step of estimations were the same for Miller et al.(2016)^30^, and a further description of full/partial correlation can be found in the document provided by UK Biobank: https://biobank.ctsu.ox.ac.uk/crystal/docs/brain_mri.pdf. Since partial correlation describes direct connectivity, this estimation was used in the present study as a major proxy for functional connectivity.

The final 55*55 and 21*21 partial correlation matrices were used as measurements of functional connections. Both types of correlation matrices were used as they addressed different principles. Since the dimensionality of the 55*55 matrix is higher, it gives a higher resolution of the whole-brain functional connectome. We therefore used this for the first two steps of whole-brain analysis. The lower dimensionality matrix, on the other hand, is better able to identify large functional networks, such as the DMN, frontal networks, and primary and higher level visual networks^23^. Hence, the functional networks that were found in the whole-brain analysis were selected from the 21*21 matrix as NOI, and the partial correlations between the NOI were used as the proxy of the coupling of these functional networks.

### Cognitive performance

A test of verbal-numerical reasoning (VNR) was carried out by UK Biobank according to standard protocol^35^. The VNR test consisted of 13 questions^24,62^. The guidelines and the questions of the test can be found in the Touch-screen fluid intelligence test protocol document: http://biobank.ctsu.ox.ac.uk/crystal/refer.cgi?id=100231). Participants were instructed to answer as quickly and correctly as possible within two minutes. The number of questions correctly answered within the time frame is the overall VNR score. The data used in the present study were collected at the time of imaging assessment (N=3,950, Age=62.07+/-7.54, Male=47.47%). The distribution of the scores in the sample analysed here is presented in Figure 1.

### Educational attainment and household income

Educational attainment and household income phenotypes were self-reported in a touchscreen-questionnaire session, the details of which are provided on the study website (http://biobank.ctsu.ox.ac.uk/crystal/refer.cgi?id=100471, http://biobank.ctsu.ox.ac.uk/crystal/refer.cgi?id=100256). Descriptive statistics of educational attainment and household income are presented in Figure 1

For educational attainment, participants could choose at least one of the following options: College or university degree, A levels/AS levels or equivalent, O levels/GCSEs or equivalent, CSEs or equivalent, NVQ or HND or HNC or equivalent, other professional qualifications, none of the above, and prefer not to answer. In accordance with the methods of previous studies^35,36^, a binary variable was created to indicate whether or not university/college level education was achieved. This proxy covered 4,160 participants (Age=62.20+/-7.56, Male=47.48%).

Household income was determined by the average total income before tax received by the participant’s household. Available choices were: <£18,000, £18,000 to £30,999, £31,000 to £51,999, £52,000 to £100,000, >£100,000, do not know and prefer not to answer. An ordinal variable from 1 to 5 was created to determine the level of household income (<£18,000 as 1, >£100,000 as 5). This measure had 3,793 non-empty responses (Age=61.98+/−7.57, Male=49.04%).

### Statistical methods

The associations between brain connections and cognitive performance, educational attainment, and household income were tested using the linear GLM function in R (https://stat.ethz.ch/R-manual/R-devel/library/stats/html/glm.html). All imaging analyses were adjusted for age, age^2^ and sex.

As the L2-regularised partial correlation represents more accurately the direct connections, as described in the section for imaging data above, we used the partial correlation matrix as the major measurement. Values in the matrix are normalised correlation coefficients. A higher absolute value means stronger strength of connection, and the sign indicates whether the connection is positive or negative. To enable clearer interpretation of the results, the values of the connections were transformed into connection strength. This was achieved by multiplying the raw connection values with the signs of their mean value. This approach was used in a previous study by Smith *et al.*(2015)^26^. For clarification purposes, the valence, mean values and 95% confidence interval of the significant connections in the section of results were reported.

Analyses were performed in the following sequence: (1) A whole-brain analysis of the association between cognitive performance (VNR) and resting-state functional connectivity. (2) Two separate whole-brain analyses on educational attainment and household income, respectively. We then performed correlation analyses on the global functional connections predicted by the three phenotypic variables, that is, testing whether the effect sizes for the VNR score’s link to functional connections was correlated with the corresponding effect sizes for educational attainment and household income. The standardised effect sizes of the non-imaging variables were extracted as a proxy of their effect on the 55*55 matrix of functional connections. Two correlation analyses were then performed respectively on (a) the effect sizes of cognitive performance and educational attainment and (b) the effect sizes of cognitive performance and household income. (3) NOI were chosen from the components from the 21-D ICA, and the tests were conducted on the associations between connections of the NOI and cognitive performance, educational attainment and household income.

False Discovery Rate (FDR)^63^ correction was applied for each model using the default settings in the p.adjust function in R (https://stat.ethz.ch/R-manual/R-devel/library/stats/html/p.adjust.html). All β-values reported in the results are standardised effect sizes.

## Acknowledgements

This study is supported by a Wellcome Trust Strategic Award “Stratifying Resilience and Depression Longitudinally” (STRADL) (Reference 104036/Z/14/Z).

This research was conducted using the UK Biobank Resource under approved project #4844. We thank the UK Biobank participants for their participation, and the UK Biobank team for their work in collecting and providing these data for analyses. Part of the work was undertaken in The University of Edinburgh Centre for Cognitive Ageing and Cognitive Epidemiology (CCACE), funding from the Biotechnology and Biological Sciences Research Council (BBSRC) and Medical Research Council (MRC) is gratefully acknowledged. Age UK (The Disconnected Mind project) also provided support for the work undertaken at CCACE.

XS receives support from China Scholarship Council. HCW is supported by a JMAS SIM fellowship from the Royal College of Physicians of Edinburgh and by an ESAT College Fellowship from the University of Edinburgh. Authors AMM, HCW, and SML gratefully acknowledge the support of the Sackler Foundation. Authors IJD, SRC and SJR are supported by the Medical Research Council award to CCACE (MR/K026992/1). IJD is additionally supported by the Dementias Platform UK (MR/L015382/1), and he and SRC by the Age UK-funded Disconnected Mind project (http://www.disconnectedmind.ed.ac.uk). SRC was supported by Medical Research Council grant MR/M013111/1.

## Author contributions

XS developed the design of the study and conducted the analyses. XS, AMM, and HCW drafted the manuscript. AMM and HCW supervised and contributed to the design of the study. IJD, SRC, SJR, DMH, SML, and MEB were involved in overseeing analysis methodology and editing the paper. MJA was involved in curating the data. UK Biobank collected all data and was involved in the preprocessing of imaging data. All authors discussed and commented on the manuscript.

